# The endosymbiont *Rickettsiella viridis* increases the virulence of *Diuraphis noxia* but reduces alate frequency

**DOI:** 10.1101/2024.09.15.613162

**Authors:** Xinyue Gu, Alex Gill, Qiong Yang, Perran A. Ross, Ella Yeatman, Mel Berran, Monica Stelmach, Sonia Sharma, Paul A. Umina, Ary A. Hoffmann

## Abstract

Aphids are among the world’s most economically damaging pests and carry a diverse range of bacterial endosymbionts. There is increasing interest in exploring potential applications of natural and novel strains of endosymbionts in aphid control. One endosymbiont, *Rickettsiella viridis,* has a large fitness cost following transfer from its natural aphid host *Acyrthosiphon pisum* into a novel host aphid *Myzus persicae*. Here, we investigated host impacts after transferring the same *Rickettsiella* strain to an important cereal aphid, the Russian wheat aphid *Diuraphis noxia*. *Rickettsiella* in this host resulted in modest fitness effects, with a minor increase in heat tolerance and a decrease in development time at 25° C. The infection persisted in mixed caged populations under different temperatures. Surprisingly, *Rickettsiella* increased aphid virulence to wheat plants and to two non-crop hosts of *D. noxia*, barley grass and brome grass. This was evident from sharper decreases in leaf number and leaf area, as well as an increase in chlorotic streaking when plants were exposed to *Rickettsiella*-infected *D. noxia*. *Rickettsiella* also reduced the proportion of alates in aphids held in small cages and in larger mesocosms containing multiple wheat plants where short-distance dispersal of aphids was impacted. These results provide compelling evidence that *Rickettsiella* can affect virulence - the first case of an endosymbiont transfer directly influencing aphid virulence to host plants – and highlight the species-specific impacts of endosymbiont transfers on aphids which can involve multiple traits.

## Introduction

Aphids (Hemiptera: Aphididae) are one of the most serious agricultural pests, damaging host plants in diverse ways (Dedryver et al., 2010) and leading to large economic losses (Valenzuela and Hoffmann, 2015, Dedryver et al., 2010, Tatchell, 1989). A substantial research effort has been directed at controlling pest aphid populations, but this remains challenging because of their high reproductive potential, ability to transmit many important plant viruses and widespread resistance to insecticides (Edwards et al., 2008, Dedryver et al., 2010). Additionally, aphids can develop winged progeny, enabling them to relocate to new host plants and disperse widely (Reyes et al., 2019).

Aphids have co-evolved with their endosymbionts for an estimated 160 to 280 million years (Fukatsu, 1994). Almost all aphids carry the obligate symbiont *Buchnera aphidicola*, which can provide essential amino acids that aphids cannot obtain from the phloem of plants (Douglas, 1998). Some aphids also carry facultative symbionts that exhibit diverse and environmentally dependent effects on hosts (Oliver et al., 2010). The impacts of endosymbionts on aphids make them a potential target for aphid control. In other insects and particularly in mosquitoes, endosymbiont manipulations are already being utilised in the field as a way to decrease viral transmission with substantial impacts on disease incidence (Hoffmann et al., 2024, Ryan et al., 2019, Utarini et al., 2021). Novel endosymbiont-related control methods are particularly promising for agricultural applications due to several factors. These include: (1) a large variety of endosymbionts in pest organisms (Sazama et al., 2019); (2) the capacity to transfer endosymbionts across species through hemolymph or cytoplasm microinjection (Chen et al., 2000, Moran and Yun, 2015, Gu et al., 2024); (3) the self-sustaining nature of endosymbionts following introduction through maternal (vertical) and/or horizontal transmission via parasitism (Gehrer and Vorburger, 2012) or plant tissue (Pons et al., 2019); (4) the large impact of endosymbionts on insect biology, including traits which can reduce plant virus transmission or pest populations (Wernegreen, 2017, Feldhaar, 2011, Cordaux et al., 2011); and (5) the species specificity of endosymbionts derived from naturally-infected insects (Eleftherianos et al., 2013).

The facultative aphid endosymbiont *Rickettsiella viridis* has been successfully transferred from its natural pea aphid host (*Acyrthosiphon pisum* (Harris)) to a novel green peach aphid host (*Myzus persicae* (Sulzer)) (Gu et al., 2023). This *Rickettsiella* infection has potential pest applications in *M. persicae* control due to its stable fitness costs (decreasing fecundity, longevity and heat tolerance) across different host plants (Ross et al., 2024), and its capacity to rapidly spread in aphid populations (Gu et al., 2023). *Rickettsiella* can also change the body color of *M. persicae* (Gu et al., 2023) and *A. pisum* (Tsuchida et al., 2010, Tsuchida et al., 2014), which could be used as a phenotypic marker and may also affect recognition by natural enemies (Soleimannejad et al., 2023). Considering its strong phenotypic effects both in its natural host *A. pisum* (Tsuchida et al., 2014, Tsuchida et al., 2010) and a novel host *M. persicae* (Gu et al., 2023), here we investigate host phenotypic effects associated with the interspecific transfer of *Rickettsiella* into the Russian wheat aphid (*Diuraphis noxia* (Mordwilko)). This aphid species is of particular interest because native bacteria present in *D. noxia* have been linked to aphid virulence in wheat (Luna et al., 2018).

*Diuraphis noxia* is a global pest of cereal crops, being found in Europe, Asia, the Americas and Australia (Ward et al., 2020). Unlike many other aphid species, *D. noxia* does not transmit plant viruses (Summers et al., 1990, Halbert et al., 1994) and rarely carries facultative endosymbionts in the wild (Luna et al., 2018, Yang et al., 2023, Ennahli et al., 2009). Economic damage through direct feeding can be substantial, particularly when *D. noxia* is at a high density (Burd and Burton, 1992, Archer and Bynum, 1992). Additionally, *D. noxia* injects a polypeptide toxin into the phloem sap while feeding (Smith et al., 2004), which affects the photosynthetic ability of the plant and prevents leaves from unrolling as they emerge (Dong et al., 1994), offering a protective leaf-roll within which the aphids reside. Because they are often hidden in rolled leaves, *D. noxia* are somewhat protected from contact insecticides and can be difficult to detect by natural enemies (Ward et al., 2020). Further complicating management efforts, *D. noxia* has a strong dispersal ability (Yazdani et al., 2018) and can persist on a large range of host plants (Hughes, 1990). *D. noxia* additionally shows chromosomal heterogeneity, allowing for greater survival of the aphid pest on host species (Nicholson et al., 2015). New biotypes are also arising, possibly due to selection pressure from resistant host cultivars (Nicholson et al., 2015), making the endosymbiotic approach attractive for pest control.

In this study, we generated a *Rickettsiella*-infected *D. noxia* strain using hemolymph microinjection and then measured its impacts on fitness, virulence to multiple host plants, alate frequency and dispersal ability. We show that the *Rickettsiella* infection can persist in caged aphid populations at different temperatures and may provide some modest fitness benefits to *D. noxia*. *Rickettsiella* also lead to an increase in aphid virulence in wheat and in two grass weeds that are known to be important non-crop hosts for this species. In addition, we show that the endosymbiont decreases the rate of alate production and dispersal within a laboratory setting.

## Materials and methods

### Aphid strains and maintenance

Aphids were cultured on Petri dishes (60 mm x 15 mm) with plant leaves placed on 1% agar. The plant host for *D. noxia* was wheat (*Triticum aestivum* cv. Trojan) and for *A. pisum* it was lucerne (*Medicago sativa*, cv. Sequel). Aphids were moved to fresh cut leaves on new Petri dishes once or twice a week and maintained in temperature-controlled cabinets at 19 (± 1) °C with a 16:8h L:D photoperiod. All plant material was grown in insect-proof BugDorm cages (90 x 45 x 45cm, mesh 160 µm aperture, Australian Entomological Supplies, Australia) within a shade-house supported with plant growth lights (40W Grow Saber LED 6500K, 1200mm length) set to a 16:8h L:D photoperiod. Wheat plants were grown for ∼ 6 weeks before use, while lucerne plants were ∼ 4.5 weeks old before use. For our experiments on virulence and aphid dispersal, wheat plants (∼ 4-6 weeks old), barley grass (*Hordeum murinum*) and brome grass (*Bromus diandrus*) (∼ 4 weeks old) were maintained at the same conditions.

A single clone of *A. pisum* as the donor species was collected from Tintinara, South Australia (GPS: -35.947 S, 140.137 E), which carried both *Serratia symbiotica* and *Rickettsiella viridis*. For the recipient *D. noxia,* we used a single clone collected from the Grains Innovation Park, Horsham (GPS: -36.723 S, 142.175E). This clone has previously been found to lack naturally occurring facultative endosymbionts (Yang et al., 2023). We generated a *D. noxia* strain carrying a stable *Rickettsiella* infection (*Rickettsiella* (+)) derived from *A. pisum* (see next section). The same clone of *D. noxia* without endosymbionts (*Rickettsiella* (-)) was used as a control in all experiments.

### Endosymbiont transfer through microinjection

We introduced *Rickettsiella* from *A. pisum* into apterous adult *D. noxia* through hemolymph microinjection (Gu et al., 2023, Chen et al., 2000). We injected 50 *D. noxia* with hemolymph from donor *A. pisum* and then followed a selection step as described in Gu et al. (2023). Endosymbiont infection status was routinely screened as well as immediately before experiments commenced through quantitative PCR (qPCR) (see “Endosymbiont detection and quantification”). Strains were approximately 40 generations post-injection before experiments commenced.

### *Effects of developmental temperature and* Rickettsiella *infection on life history*

Two nymphs (< 24 h old) were transferred to a plastic cup (150 ml: 8 x 6 x 5 cm, Daily Best, Auburn, Australia) with an individual wheat plant (root removed) inserted into 2 cm 1% agar. A second cup was placed on top and sealed with tape. Thirty replicates each were set up for *Rickettsiella* (-) and *Rickettsiella* (+) *D. noxia* at each temperature of 19°C (± 1°C) and 25°C (± 1°C) under plant growth lights. After 7 d, a single aphid was randomly removed and stored in 100% ethanol for endosymbiont density measurements. Aphids were transferred to new cups every week. Developmental time, lifetime fecundity and longevity were monitored as described in Gu et al. (2023).

### Body color and body shape

Unlike in *M. persicae* (Gu et al., 2023) and *A. pisum* (Tsuchida et al., 2010), there was no obvious effect of *Rickettsiella* on body color after transfer. Nevertheless, to test for subtle differences, adult apterous aphids (∼ 14 d old) from 19°C were assessed in detail, with 25 individuals from each of the *Rickettsiella* (-) and *Rickettsiella* (+) strains measured. We measured body length, body width and body color by taking photos and analyzing these with ImageJ, as described previously (Gu et al., 2023).

### CTmax and knockdown time for heat tolerance

We measured the heat tolerance using CTmax estimated from heat knockdown assays (Bak et al., 2020) as described in Gu et al. (2023). Seven-day old aphids were used in both assays, with the temperature in CTmax increased from 25°C at a rate of 0.1°C per min, and knockdown time set to a constant temperature of 42°C until all aphids became incapacitated. We performed two blocks for both assays, with 30 individuals from each strain measured in each block.

### Rickettsiella *population dynamics*

Thirty *Rickettsiella* (+) and 30 *Rickettsiella* (-) mixed aged *D. noxia* were transferred to the leaves of six wheat plants (∼ 6 weeks old) and placed in insect proof BugDorm cages (30 x 30 x 62 cm, mesh 160 µm aperture, Australian Entomological Supplies, Australia). Each cage was replicated five times for each temperature of 19°C and 25°C. Aphids were transferred to new wheat plants every 3 weeks at 19°C and every 2 weeks at 25°C. At each transfer, sixty randomly selected aphids were placed on the new plants, with the remaining aphids stored in 100% ethanol. To measure *Rickettsiella* frequency over time, we screened 15 aphids per replicate cage at multiple time points (Weeks 3, 9 and 12 at 19°C and weeks 2, 8 and 12 at 25° C) using qPCR.

### Virulence to wheat plants

As *D. noxia* causes yield loss by direct feeding and chlorosis of leaves by injecting toxins into the plants (Damsteegt et al., 1992), we compared the virulence of *Rickettsiella* (+) and *Rickettsiella* (-) aphids to individual wheat plants. For this comparison, two development stages of wheat plants (LongReach Plant Breeders Management Pty Ltd, Australia) were used: Zadoks GS39 (∼ 6 weeks old), with plants at flag leaf emergence; and Zadoks GS13 (∼ 4 weeks old), where plants had three unfolded leaves.

For the experiment involving wheat plants at GS39, ten mixed-aged apterous adults from each strain were placed onto second and third true leaves of a single plant growing in a plastic gardening pot containing potting mix. Perforated bread stick plastic bags and rubber bands were used to seal each plant, which was then randomly placed at 25°C (± 1°C) with a 16:8 h L:D photoperiod. Ten replicates were established for each treatment: *Rickettsiella* (+), *Rickettsiella* (-), and wheat plants with no aphids added. Survival of aphids was assessed after 24 h to ensure all individuals remained alive. Each week, we recorded the total leaf number, leaf area (summed across leaves as width*length*0.75 (Aldesuquy et al., 2014)), the percentage of leaves with chlorotic streaking and aphid colonization rate (the percentage of leaves with more than ten adults); the first two were measured from the week 2 onwards after aphids had colonized the plants. The experiment was terminated when plants in one of the replicates had mostly died, which was at week 4.5. The above-ground plant matter (minus the wheat heads) within each replicate was removed, dried at 60 °C for 24 h, and then weighed to estimate plant biomass.

For the experiment involving wheat plants at GS13, five age-matched apterous adults (14 d old) from each strain were placed onto the second true leaf of a single wheat plant. Survival of aphids was again assessed after 24 h. Perforated bread stick plastic bags and rubber bands were used to seal each plant, which was randomly placed at 25°C (± 1°C) with a 16:8 h L:D photoperiod. Sixteen replicates were established for each treatment: *Rickettsiella* (+), *Rickettsiella* (-), and wheat plants with no aphids added. Each week, we recorded the total leaf number, leaf area, the percentage of leaves with chlorotic streaking and aphid colonization rate as described above. Additionally, we measured a weekly aphid infestation score and overall plant damage score for all replicates (see Table 1). All scores were assessed independently by three people and these were later averaged to provide overall scores at each time point. After the week 2 assessment, we removed three replicates with individual wheat plant as a single replicate from each treatment to characterize the developmental stage of aphids every week. Here, we recorded the aphid morph and life-stage (alate adults, apterous adults and nymphs). The experiment was terminated when plants in one of the replicates had a plant damage score of 4 or above, which was at week 3.5. The above-ground plant matter within each replicate was then removed, dried at 60 °C for 24 h, and weighed.

**Table 1.**
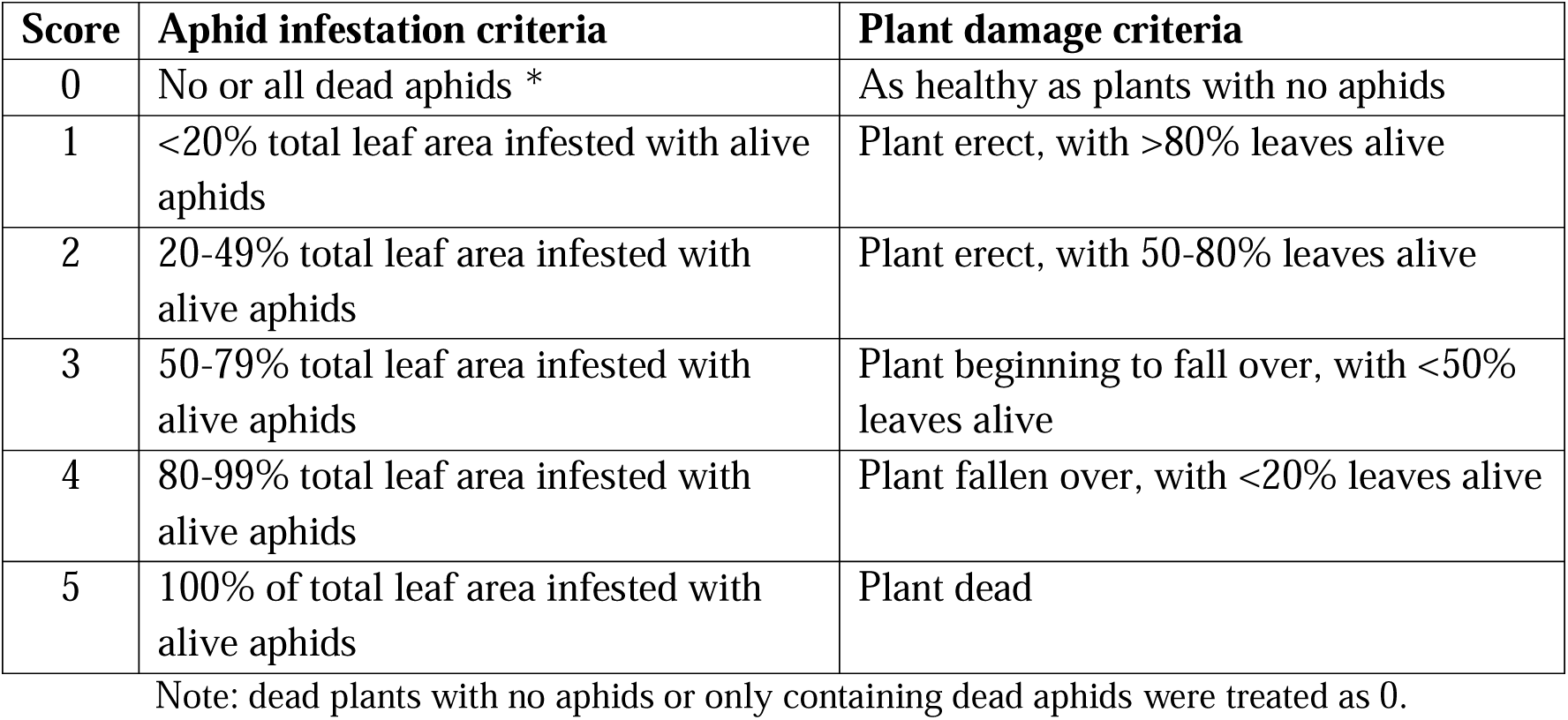
Criteria for aphid infestation and plant damage scores.

### Virulence to non-crop plants

Two grass weeds that are known plant hosts for *D. noxia*, barley grass (*Hordeum murinum*) and brome grass (*Bromus diandrus*), were assessed for virulence of *Rickettsiella* (+) and *Rickettsiella* (-) strains. Wheat plants were used as a control.

To control for plant age, wheat, barley grass and brome grass were germinated in Petri dishes covered by wet filter paper, with seedlings subsequently transplanted into plastic gardening pots containing potting mix. Plants were then placed into a temperature-controlled cabinet with plant growth lights (TS1000 150W Full Spectrum LED Grow Light, Mars Hydro, Australia; dimmed to 50%) at 20°C (± 1.5°C) with a 16:8h L:D photoperiod. The plants were used for the experiments 4 weeks later. Each replicate consisted of two plants of a species placed in a BugDorm cage (30 x 30 x 62 cm). Five replicates for each strain/plant species combination were randomly placed at 25°C with a 16:8h L:D photoperiod under plant growth lights (40W Grow Saber LED 6500K, 1200mm length). Based on plant size, three age-matched apterous adults (14 d old) from each strain were placed into each pot containing wheat plants and brome grass plants, or two 14 d old aphids for each pot containing barley grass. After 24 h, aphid survival was assessed. Once or twice per week, we recorded the total leaf number, leaf area (after week 2), chlorotic streaking, aphid colonization rate, aphid infestation score and plant damage score, as described above. The experiment was terminated when most plants within one replicate were given a plant damage score of 4 or above, which was at week 2.5 for wheat plants, week 3 for barley grass and week 5 for brome grass. All aphids from each replicate were separated into apterous adults, alate adults and nymphs, and counted.

### Aphid dispersal ability and virulence to wheat in mesocosms

As the number of alate and apterous adults in the virulence microcosm experiment varied among strains, which might influence aphid dispersal ability, a larger scale experiment was undertaken. Three plastic trays (39.5 x 28 x 11cm, Bargain Basement, Australia) with eight wheat plants (GS13) per tray were established. These were placed inside a single insect proof BugDorm cage (90 x 45 x 45cm, mesh 160 µm aperture) as a replicate, creating a mesocosm environment. Five replicate mesocosms were established and randomly positioned in a temperature-controlled room at 25 (± 1) °C with a 16:8 h L:D photoperiod. For each replicate, *Rickettsiella* (+) and *Rickettsiella* (-) aphids were released at either end of the BugDorm cage, with 5 age-matched individuals (14 d old) placed on the second true leaf of each plant (n= 4 wheat plants each side). Aphid survival was assessed after 24 h. The four plants with aphids represented the ‘aphid-release wheat’ and the four adjacent plants represented the ‘aphid-spread wheat’. The middle tray with wheat plants was used to further monitor aphid dispersal. Once or twice per week, we recorded the total leaf number, leaf area, chlorotic streaking, aphid colonization rate, aphid infestation score and plant damage score, as described above. To assess aphid dispersal, all eight wheat plants in the middle tray were removed once per week and replaced with fresh plants at the same growth stage. All aphids on the eight plants were subsequently removed, counted and stored in 100% ethanol at -20°C. In addition, 20 alates and 20 apterous aphids at each sampling point within each mesocosm were randomly selected to screen for endosymbiont infection status.

### Endosymbiont detection and quantification

qPCR assays were used to confirm the presence or absence of *Rickettsiella* and/ or *Buchnera* infection and to measure their density relative to a host gene following the methods of Lee et al. (2012). Relative *Rickettsiella* and *Buchnera* densities were determined as described in Gu et al. (2023) and then transformed by 2^ΔCp^. Units presented in the figures are the transformed values shown on a log scale. In the experiments assessing life history traits and population dynamics, aphids with a *Rickettsiella* Cp value <30 and Tm values within the range of positive controls (87.5-88.3) considered *Rickettsiella* positive and high density, while aphids with a *Rickettsiella* Cp value >30 were considered *Rickettsiella* positive but with low density, and negative if Cp values were absent and Tm values were not within the range of positive controls. In the mesocosm experiment assessing aphid dispersal and virulence, aphids with a *Rickettsiella* Cp value >30 were considered *Rickettsiella* negative due to the potential for horizontally transmitted *Rickettsiella* to lead to *Rickettsiella* (-) aphids testing positive with low densities (Figure S1).

### Statistical analysis

Phenotypic data were analysed with IBM SPSS Statistics 29.0 for Mac following normality tests. Development time measures were analysed with Kruskal-Wallis Tests. Fecundity was analysed with general linear models (GLMs), with developmental temperature, *Rickettsiella* infection status and their interaction included as factors. Endosymbiont density values were compared by independent sample t-tests. We used Cox regression to assess the impact of *Rickettsiella* infection on aphid longevity. For body color and body shape, we analysed each component (hue, saturation, lightness) separately using independent sample t-tests as well as all the components together as a multivariate ANOVA (MANOVA). For thermal tolerance traits (heat knockdown time and CTmax), we included block as a random factor and *Rickettsiella* infection status as a fixed factor and performed analyses using GLMs. For the population dynamics experiment, GLMs were used to test for changes in *Rickettsiella* infection status across time. For the virulence experiments, we focused on the difference between *Rickettsiella* (+) and *Rickettsiella* (-) aphid-infested plants using independent t-tests at each sampling point, as well as repeated measures in GLMs for the wheat GS39 experiment, non-crop experiment and the mesocosm wheat experiment. Additionally, we performed a MANOVA with a combination of all plant traits to test the overall effects of *Rickettsiella* (+) and *Rickettsiella* (-) in all virulence experiments. A one-way ANOVA was used to analyse the plant biomass data. Aphid dispersal data from the mesocosm experiment was analysed by paired t-tests.

## Results

### Rickettsiella *transfer and life history impacts*

We successfully transferred *Rickettsiella* from the donor *A. pisum* to *D. noxia* through hemolymph microinjection. Of the 50 aphids injected, 6/12 surviving aphids produced nymphs that tested positive for *Rickettsiella* and one of these passed *Rickettsiella* to its offspring stably. Although *A. pisum* carried both *Serratia symbiotica* and *Rickettsiella* infections, the *Serratia* infection was not detected in *D. noxia* at G0, while the *Rickettsiella* infection was stable at a frequency of 100% for over 50 generations following selection.

We measured the effects of *Rickettsiella* in *D. noxia* on life history traits under two different temperatures (19 and 25 °C), as well as body color and body shape at 19 °C (Figure 1). There was a significant effect of *Rickettsiella* on development time (Kruskal-Wallis Test: H_1,62_

**Figure 1.**
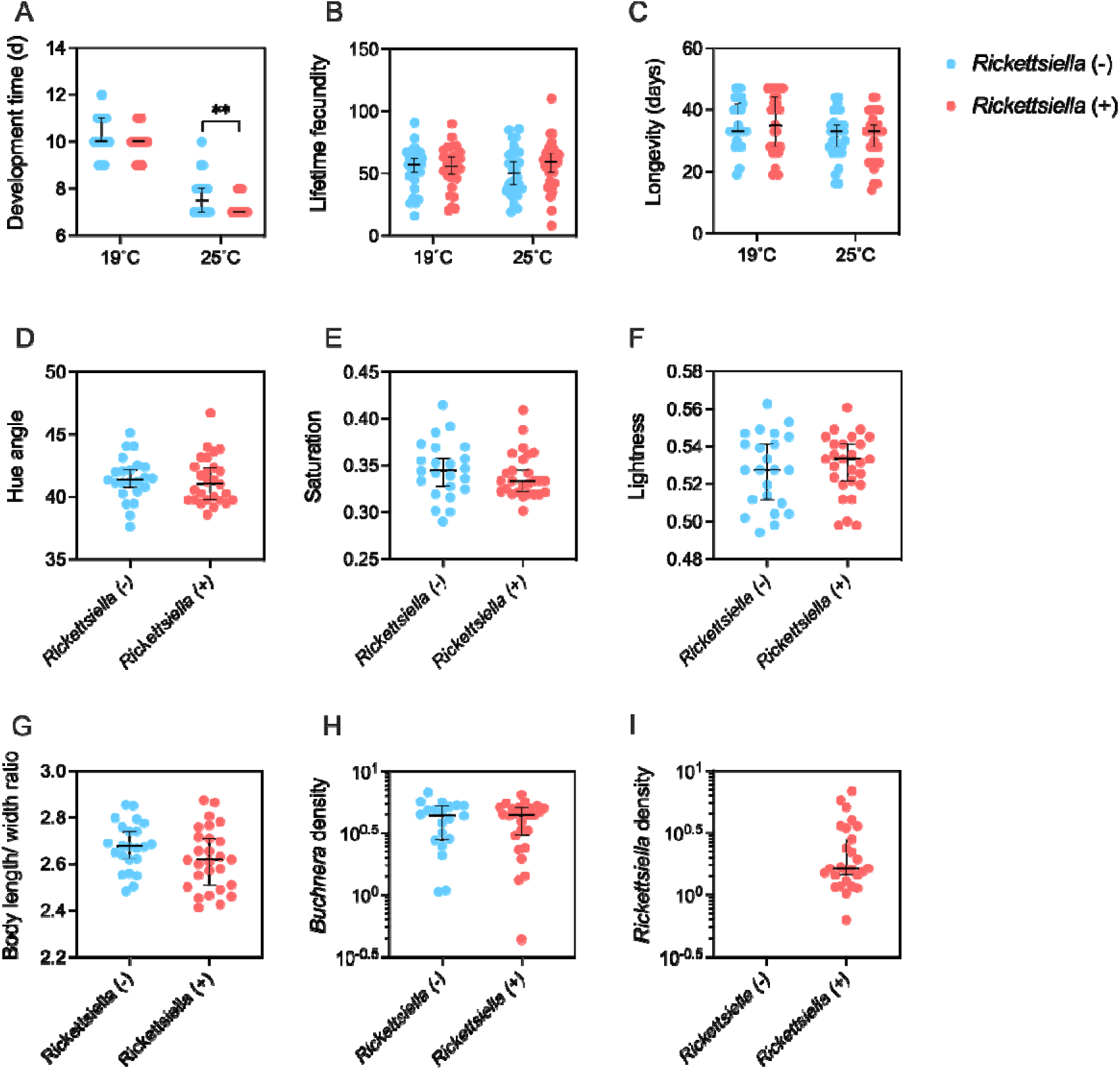
Effects of *Rickettsiella* on the life history, body color, body shape and *Buchnera* density of *D. noxia*. Aphids from *Rickettsiella* (+) and *Rickettsiella* (-) reared at 19 °C and 25 °C were individually measured for life history parameters: (A) development time, (B) lifetime fecundity, and (C) longevity. Aphids reared at 19°C in groups were measured for (D-F) body color and (G) body shape (expressed as the ratio of body length/body width). We also quantified (H) *Buchnera* and (I) *Rickettsiella* density relative to actin, where numbers represent the average difference in Cp values between endosymbiont and actin markers, transformed by 2^ΔCp^. Body color was separated into three components: (D) Hue, (E) Saturation and (F) Lightness. Dots represent data from individual aphids, while horizontal black lines depict medians. Error bars indicate 95% confidence intervals.

=4.405, *P* = 0.036, Figure 1A), with *Rickettsiella* (+) aphids having a faster development time compared with *Rickettsiella* (-) aphids at 25 °C. However, there was no significant difference in lifetime fecundity (GLM: F_1,118_ = 1.321, *P* = 0.253, Figure 1B) or longevity (Cox regression: χ^2^ = 0.153, d.f. = 1, *P* = 0.696, Figure 1C) between *Rickettsiella* (+) and *Rickettsiella* (-) aphids. We found a significant effect of temperature on *D. noxia* life history traits, except for fecundity (Kruskal-Wallis test: development time: H_1,122_ =86.061, *P* < 0.001; longevity: Cox regression: χ^2^ = 10.635, d.f. = 1, *P* = 0.001; GLM: lifetime fecundity: F_1,118_ = 0.033, *P* = 0.857). Unlike in *M. persicae* (Gu et al., 2023)*, Rickettsiella* infection had no obvious effect on body color in *D. noxia* (MANOVA across all color components: *P* =0.618; t-test: Hue: *P* = 0.795, Figure 1D; Saturation: *P* = 0.409, Figure 1E; Lightness: *P* = 0.603, Figure 1F). The body shape of *D. noxia* apterous adults was also not influenced by *Rickettsiella* infection (t-test: *P* = 0.115, Figure 1G). Separate data for body length was also non-significant (*P* =0.092, Figure S2A), while *Rickettsiella* (+) aphids were slightly wider than *Rickettsiella* (-) aphids (*P* = 0.013, Figure S2B).

We measured *Buchnera* and *Rickettsiella* densities at 19 °C and found no effect of *Rickettsiella* infection on *Buchnera* density (t-test: *P* = 0.914, Figure 1H), suggesting the addition of a novel *Rickettsiella* infection did not perturb the primary endosymbiont *Buchnera*. All *Rickettsiella* (+) aphids we screened were positive for *Rickettsiella* infection, with a median density of around 10^0^ (*i.e.,* one copy of the endosymbiont per host control gene) (Figure 1I), while all *Rickettsiella* (-) aphids screened were negative.

### Rickettsiella *infection increases aphid heat tolerance*

We tested the effect of *Rickettsiella* on *D. noxia* heat tolerance by estimating CTmax and heat knockdown time (Figure 2). For CTmax, *Rickettsiella* infection (GLM: F_1,100_ = 3.408, *P* = 0.316) and block (F_1,100_ = 2.643, *P* = 0.351) had no significant effects overall (Figure 2A), although when data were analyzed separately, *Rickettsiella* infection increased CTmax in both blocks (t-test: all *P* < 0.05). In the heat knockdown experiment, *Rickettsiella* (+) aphids showed a significantly longer time to knock down compared with *Rickettsiella* (-) aphids (GLM: F_1,109_ = 380236.17, *P* = 0.001, Figure 2A). We also found a significant effect of block (F_1,108_ = 11407.19, *P* = 0.006), although *Rickettsiella* infection significantly reduced knockdown time in both blocks (t-test: all *P* < 0.001).

**Figure 2.**
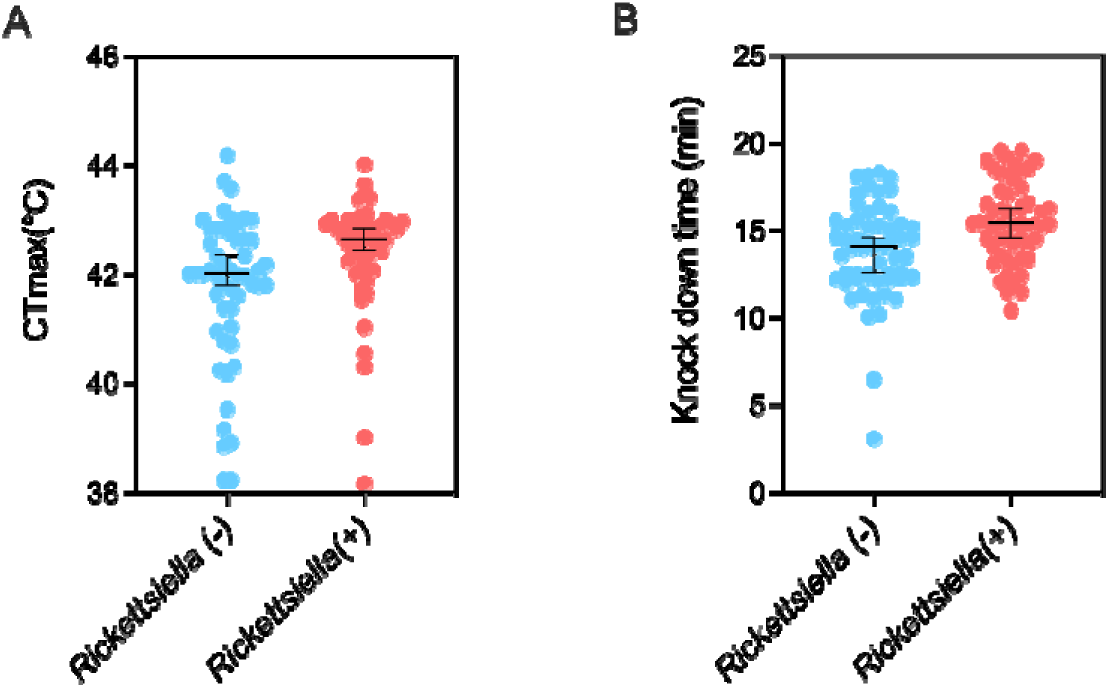
Effects of *Rickettsiella* infection on *D. noxia* heat tolerance. *Rickettsiella* (+) and *Rickettsiella* (-) aphids were measured for (A) CTmax at a ramping rate of 0.1°C/min from 25 °C, and (B) heat knockdown time at a constant 42.5°C. Dots represent data from individual aphids, while black horizontal lines are medians. Error bars indicate 95% confidence intervals. The data are shown with the blocks combined, while data for each separate block is presented in Figure S3.

### *Persistence of* Rickettsiella *in BugDorm cages depends on temperature*

We established five BugDorm cages at 19 °C and 25 °C with mixed populations of *Rickettsiella* (+) and *Rickettsiella* (-) aphids to investigate population dynamics across time (Figure 3A). The proportion of *D. noxia* positive for *Rickettsiella* increased rapidly, with a frequency above 75% in most of replicates by week 2 at 25 °C (Figure 3C) and week 3 at 19 °C (Figure 3B). At 25°C, *Rickettsiella* showed a high prevalence until week 12 (Figure 3C) but at 19°C, there was a decline from week 9 to 12 (Figure 3B). Horizontal transmission is usually expected to produce aphids with a low density of the endosymbiont initially, which then potentially increases and is passed down to ensuing generations over time (Gu et al., 2023). Aphids with a low density of *Rickettsiella* were found to be common but were seen at a relatively low frequency at 25 °C (Figure 3G). Aphids with a high density of *Rickettsiella* typically kept stable over time at 25 °C (Figure 3E). However, aphids with a high density of *Rickettsiella* persisted at similar levels to aphids with low *Rickettsiella* density at 19 °C (Figures 3D & 3F). Overall, *Rickettsiella* density of infected *D. noxia* remained relatively stable across time at both 19 °C and 25 °C, with no effect of sampling date (GLM: F_1,326_ =0.279, *P* = 0.598, Figure S4) or temperature (F_1,326_ = 3.649, *P* = 0.128, Figure S4).

**Figure 3.**
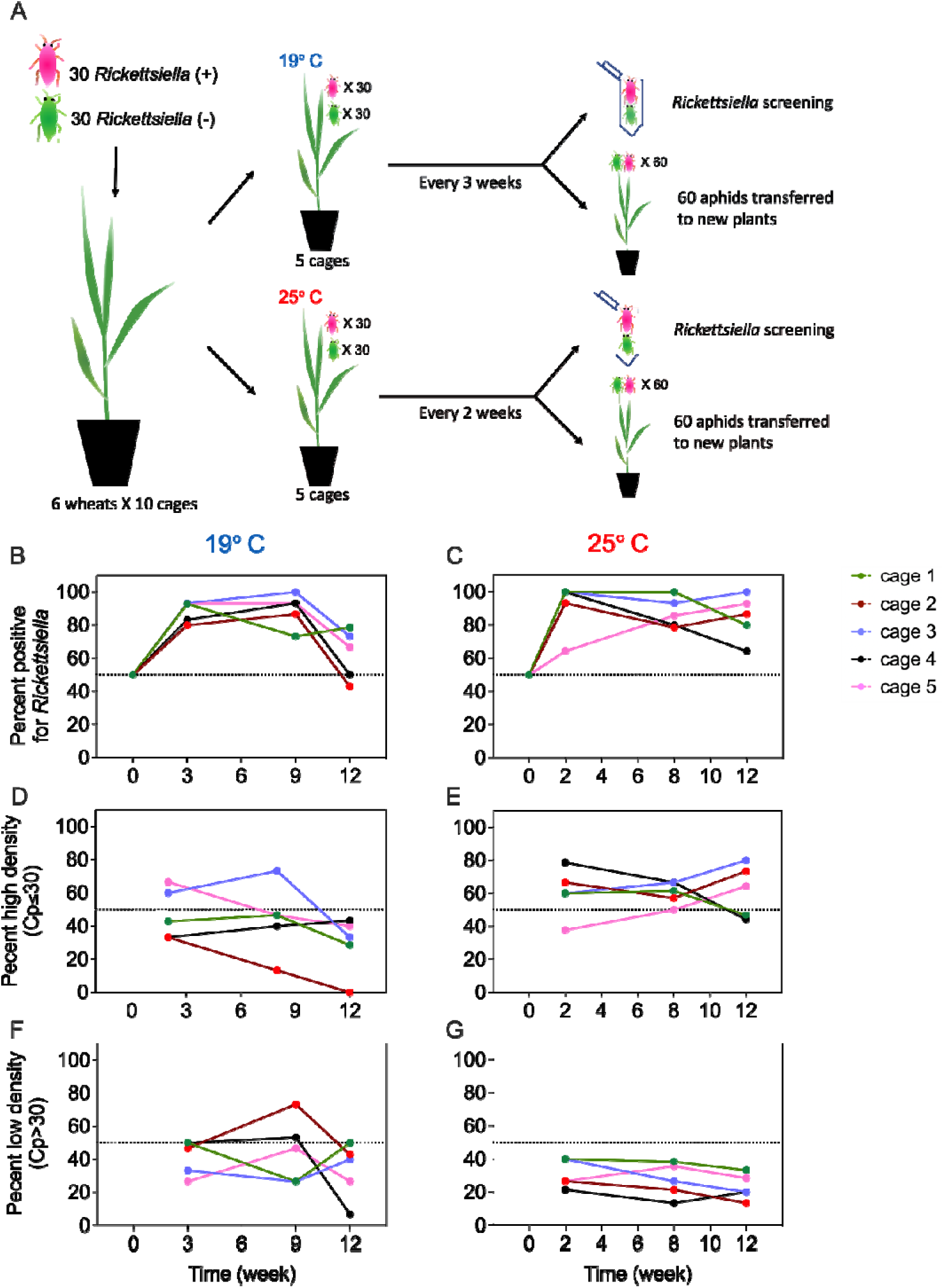
Population dynamics of *Rickettsiella* in BugDorm cages at 19 °C and 25 °C. (A) Experimental design. Populations were initiated with 30 *Rickettsiella* (+) and 30 *Rickettsiella* (-) *D. noxia* on wheat plants placed in BugDorm cages. Five replicates were set up with six wheat plants per cage and maintained at either 19° C or 25° C. Samples of 60 aphids were transferred to new plants every 3 weeks at 19 °C and every 2 weeks at 25 °C. The remaining aphids were stored for *Rickettsiella* infection dynamics at (B) 19 °C and (C) 25 °C. We tracked the *Rickettsiella* infection rate for individuals with a high density of the endosymbiont (with Cp<=30) and low density (Cp>30) of the endosymbiont at (D, F) 19 °C and (E, G) 25 °C. Aphids were considered negative if Cp values were absent and Tm values were not within the range of positive controls (87.5-88.3). Symbols represent the proportion of individuals testing positive for *Rickettsiella* at 15 aphids per time point, per replicate cage. Infection density data are shown in Figure S4.

### Rickettsiella *infection increases aphid virulence to wheat plants and reduces alate production*

We established an experiment to measure the impact of aphids with or without *Rickettsiella* on plant health. Compared with healthy un-infested wheat plants, the presence of *D. noxia* led to severe physical damage (Figures 4 & S5). Moreover, *Rickettsiella* infection increased aphid virulence, regardless of plant development stage.

**Figure 4.**
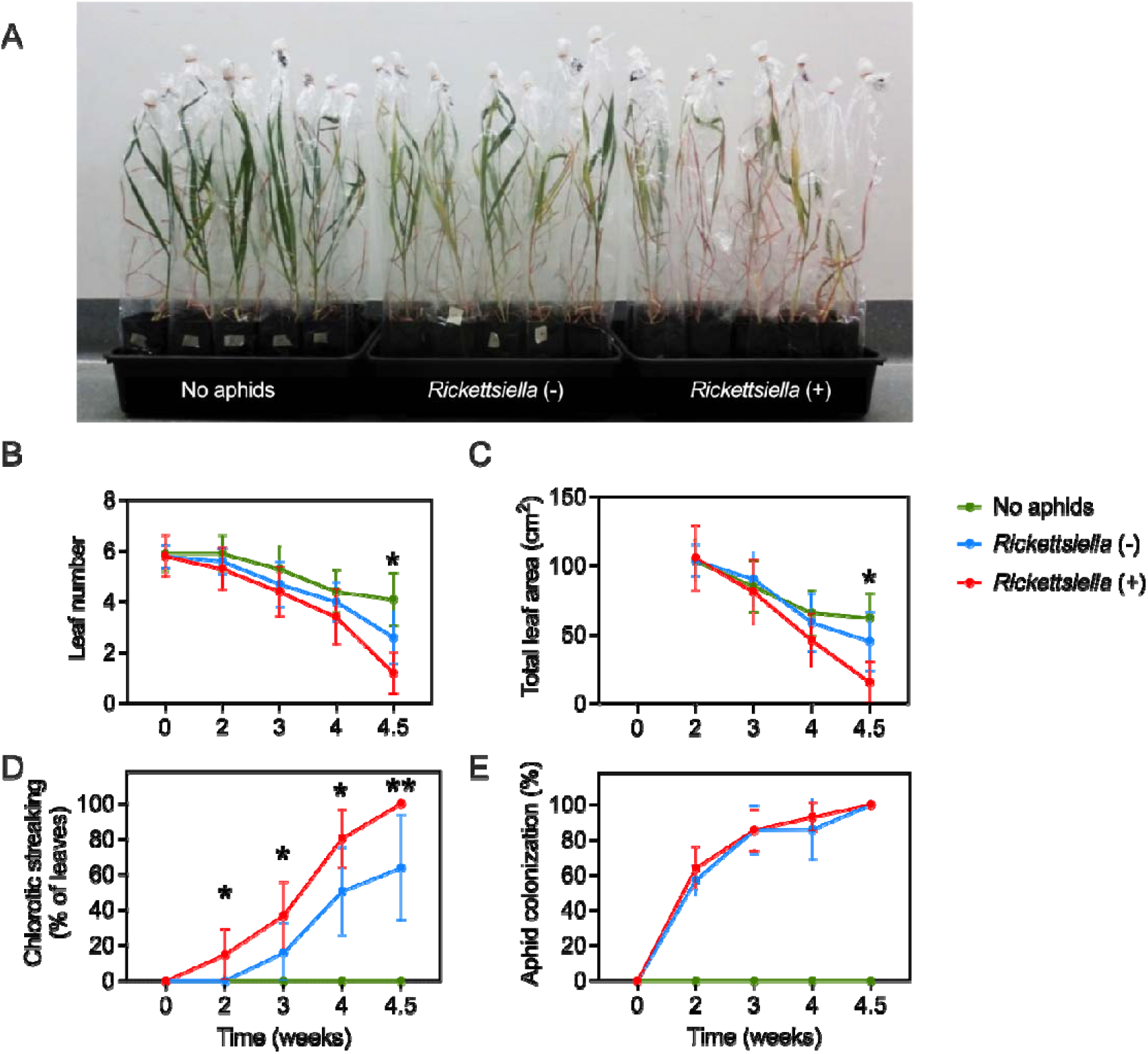
Aphid virulence to wheat plants at GS39 when infested with *Rickettsiella* (+) or *Rickettsiella* (-) *D. noxia*. Ten mixed aged *Rickettsiella* (+) or *Rickettsiella* (-) *D. noxia* were placed on individual wheat plants. Clean wheat plants (without any aphids) were included for comparison. (A) Aphid virulence to wheat plants when infested with *Rickettsiella* (+), *Rickettsiella* (-) *D. noxia* or when lacking aphids at week 4.5. Each treatment had ten replicates and measured plant traits included (B) leaf number, (C) total leaf area, (D) chlorotic streaking (expressed as a percentage) and (E) aphid colonization rate. “*” and “**” represent significant differences at *P* < 0.05 and *P* < 0.01, respectively comparing *Rickettsiella* (+) and *Rickettsiella* (-) treatments by independent sample t tests.

When plants were infested at GS39, aphids caused leaves to rapidly die (Figure 4A). *Rickettsiella* (+) aphids produced significantly more damage at week 4.5 compared with the *Rickettsiella* (-) aphids, with plants having fewer leaves (t-test: *P* = 0.026; Figure 4B) and a decrease in total leaf area (t-test: *P* = 0.018; Figure 4C). The levels of chlorotic streaking, a typical symptom of *D. noxia* feeding, were always greater and were expressed earlier in plants infested with *Rickettsiella* (+) aphids compared with *Rickettsiella* (-) aphids (t-test: all *P* < 0.05; Figure 4D). This is despite the overall aphid colonization being very similar between treatments (Figure 4E). When considering the repeated measurements across time, wheat plants infested with *Rickettsiella* (+) aphids had higher chlorotic streaking (GLM: F_1,17_ = 19.233, *P* < 0.001), and fewer (but non-significant) leaves (F_1,17_ = 3.975, *P* = 0.062) and leaf area (F_1,17_ = 3.337, *P* = 0.085). For plant biomass, treatment effects were not significant (*P* = 0.374), even when un-infested plants were included (GLM: F_2, 27_ = 2.713, *P* = 0.085, Figure S5A) or when using a MANOVA (*P* = 0.279).

When infested by *D. noxia* at GS13, wheat plants died more quickly, and aphids showed increased virulence across time. At week 3, we saw the largest difference between plants infested with *Rickettsiella* (+) and *Rickettsiella* (-) aphids, with fewer leaves (t-test: *P* = 0.041; Figure 5A) and a higher frequency of chlorotic streaking (t-test: *P* = 0.005; Figure 5C) in the *Rickettsiella* (+) treatment. There were no significant differences in total leaf area (Figure 5B), aphid colonization (Figure 5D), aphid infestation score (Figure 5E) or overall plant damage score (Figure 5F) between *Rickettsiella* treatments. Additionally, regardless of plant trait considered, from week 2 to 3.5, the level of virulence did not significantly differ between treatments or when applying a MANOVA (all: *P* > 0.05). Early aphid infestation had a large influence on plant biomass (GLM: F_2, 27_ = 47.90, *P* < 0.001, Figure S5B) but no difference was found between the *Rickettsiella* (+) and *Rickettsiella* (-) treatments (*P* > 0.05).

**Figure 5.**
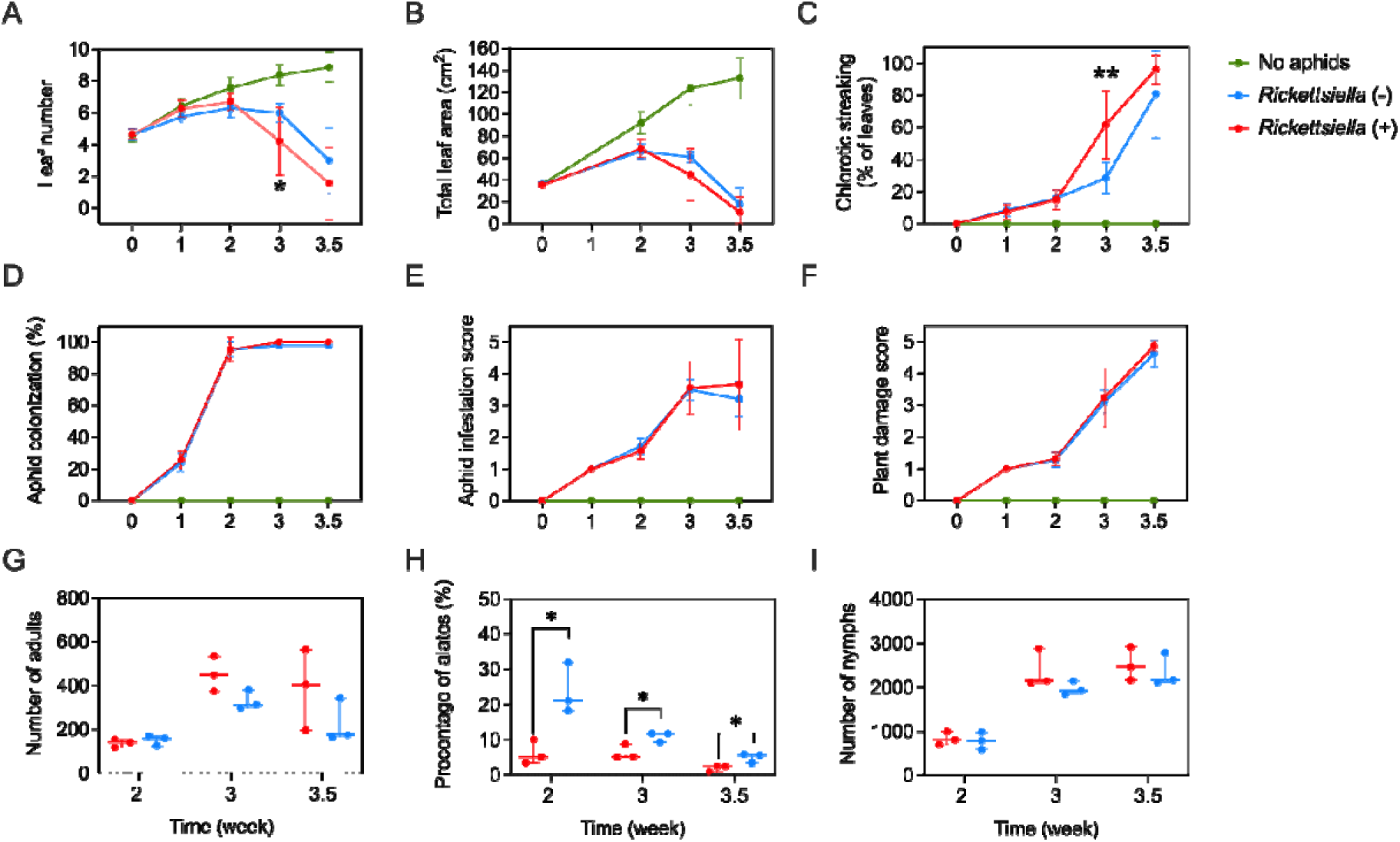
Aphid virulence to wheat plants at GS13 when infested with *Rickettsiella* (+) or *Rickettsiella* (-) *D. noxia* and effects on aphid developmental stage. Five age-matched *Rickettsiella* (+) or *Rickettsiella* (-) *D. noxia* were placed on wheat plants. Clean wheat plants without aphids were included for comparison. Each treatment had 16 replicates and measured traits were (A) leaf number, (B) total leaf area, (C) chlorotic streaking, (D) aphid colonization rate, as well as (E) an aphid infestation score and (F) plant damage score. From week 2 onwards, aphids from a single replicate *Rickettsiella* (+) and *Rickettsiella* (-) treatment (three replicates were used per week) were removed and the number of alate adults, apterous adults and nymphs recorded. (G) The number of adults (alate plus apterous adults), (H) the percentage of alates and (I) the number of nymphs were used to characterize the developmental stages of aphids over time. “*” and “**” represent significant differences at *P* < 0.05 and *P* < 0.01, respectively comparing *Rickettsiella* (+) and *Rickettsiella* (-) treatments by independent sample t-tests.

When considering developmental stages, we found the number of alate adults, apterous adults and nymphs changed significantly over time in both *Rickettsiella* (+) aphids (GLM: all *P<* 0.045) and *Rickettsiella* (-) aphids (GLM: all *P* < 0.032). Moreover, there were significantly fewer alates as a percentage from week 2 to week 3.5 (t-test: all *P* < 0.034, Figure 5H) when there were similar total numbers of *Rickettsiella* (+) and *Rickettsiella* (-) adults and nymphs on plants (t-test: all *P* < 0.08, Figures 5G & 5I). The percentage of alates decreased as the plants became more damaged (Figure 5H) but remained significantly lower for the *Rickettsiella* (+) strain at all sampling points.

### Rickettsiella *infection increases aphid virulence to non-crop plants*

The increased virulence of the *Rickettsiella* (+) strain observed in wheat plants was also evident in two non-crop plants, barley grass and brome grass. For barley grass, plants infested with *Rickettsiella* (+) aphids died much quicker (Figure 6A), with fewer leaves (t-test, *P* = 0.044, Figure 6C), lower total leaf area (*P* = 0.046, Figure 6D), higher chlorotic streaking (*P* < 0.001, Figure 6E) and more severe plant damage (*P* = 0.037, Figure 6G) compared with plants exposed to *Rickettsiella* (-) aphids. When a combination of all plant traits was considered in the MANOVAs, virulence to barley grass differed significantly between aphid strains at week 3 (*P* = 0.040). When considering the repeated measurements across time from week 1.5 to week 3, barley grass infested with *Rickettsiella* (+) aphids consistently had a higher chlorotic (GLM: F_1,18_ = 22.281, *P* < 0.001).

**Figure 6.**
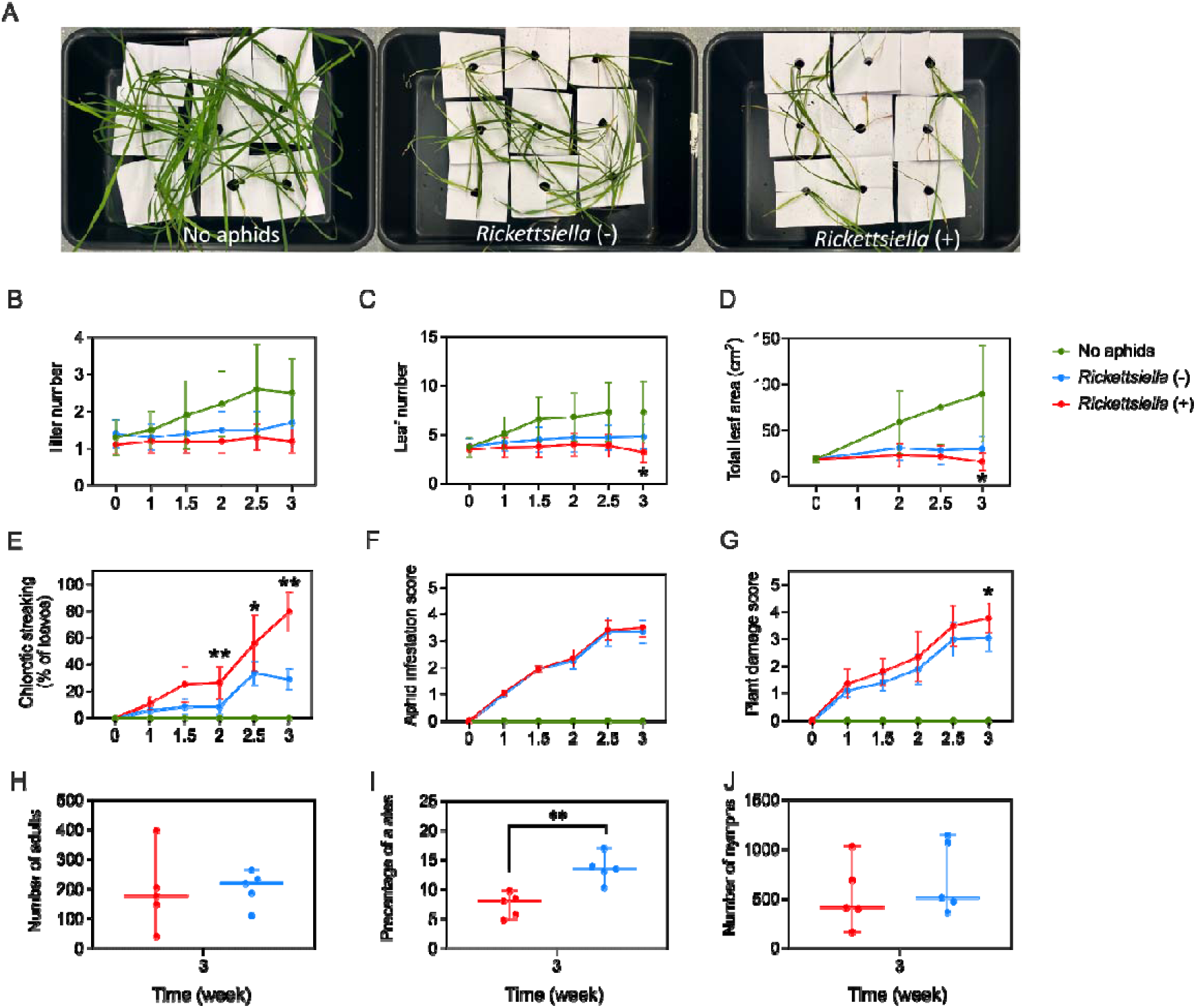
Aphid virulence to barley grass infested with *Rickettsiella* (+) or *Rickettsiella* (-) *D. noxia* and effects on aphid developmental stage. Two age-matched *Rickettsiella* (+) or *Rickettsiella* (-) *D. noxia* were placed on individual barley grass plants. Clean plants without aphids were included for comparison. (A) Aphid virulence to barley grass when infested with *Rickettsiella* (+), *Rickettsiella* (-) *D. noxia* or when lacking aphids at week 3. Each treatment had 5 replicates with 2 plants per replicate and measured traits were (B) tiller number, (C) leaf number, (D) total leaf area, (E) chlorotic streaking, as well as (F) an aphid infestation score and (G) plant damage score. At week 3, aphids from the *Rickettsiella* (+) and *Rickettsiella* (-) treatments were removed and the number of alate adults, apterous adults and nymphs were recorded. (H) The number of adults (alate plus apterous adults), (I) the percentage of alates and (J) the number of nymphs were used to characterize the developmental stages of aphids. “*” and “**” represent significant differences at *P* < 0.05 and *P* < 0.01, respectively comparing *Rickettsiella* (+) and *Rickettsiella* (-) treatments by independent sample t-tests.

For brome grass, most plant traits were similar between aphid strains (Figure 7) except for chlorotic streaking, where *Rickettsiella* (+) aphid-infested plants consistently had higher streaking across time based on repeated measurements (GLM: F_1,18_ = 5.488, *P* = 0.031).

**Figure 7.**
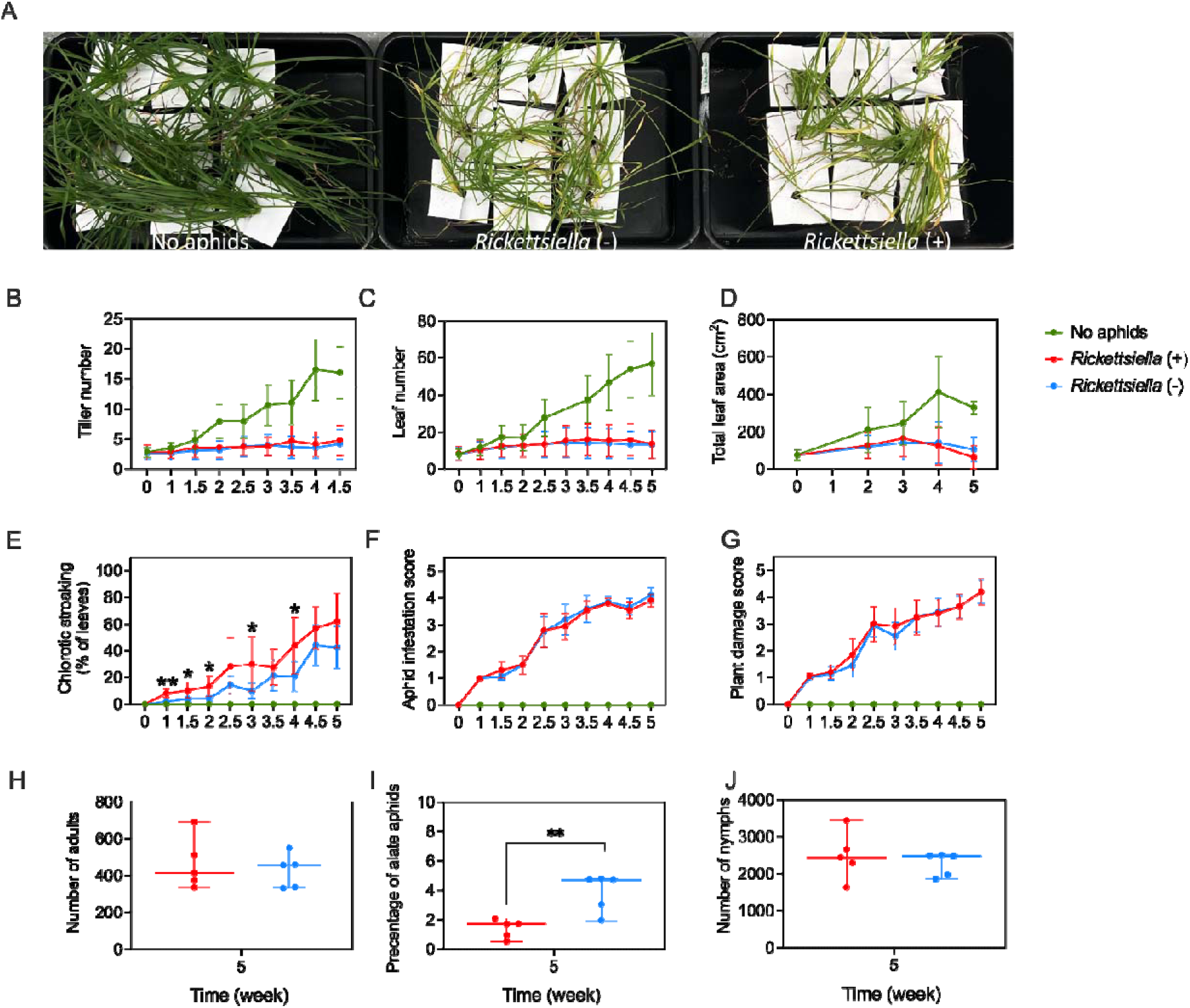
Aphid virulence to brome grass when infested with *Rickettsiella* (+) or *Rickettsiella* (-) *D. noxia* and effects on aphid developmental stage. Three age-matched *Rickettsiella* (+) or *Rickettsiella* (-) *D. noxia* were placed on individual brome grass plants. Clean plants without aphids were included for comparison. (A) Aphid virulence to brome grass when infested with *Rickettsiella* (+), *Rickettsiella* (-) *D. noxia* or when lacking aphids at week 5. Each treatment had 5 replicates with 2 plants per replicate and measured traits were (B) tiller number, (C) leaf number, (D) total leaf area, (E) chlorotic streaking, as well as (F) an aphid infestation score and (G) plant damage score (G). At week 5, aphids from the *Rickettsiella* (+) and *Rickettsiella* (-) treatments were removed and the number of alate adults, apterous adults and nymphs recorded. (H) The number of adults (alate plus apterous adults), (I) the percentage of alates and (J) the number of nymphs were used to characterize the developmental stages of aphids. “*” and “**” represent significant differences at *P* < 0.05 and *P* < 0.01, respectively comparing *Rickettsiella* (+) and *Rickettsiella* (-) treatments by independent sample t-test.

The effects on wheat were similar to our earlier experiment assessing virulence to this plant. A higher level of chlorotic streaking was observed in the *Rickettsiella* (+) treatment compared with the *Rickettsiella* (-) treatment from week 2 onwards (Figure S6).

When considering developmental stages, *Rickettsiella* (+) aphids showed a lower alate frequency compared with the *Rickettsiella* (-) aphids on all three plant species (barley grass: *P* = 0.002, Figure 6I; brome grass: *P* = 0.005, Figure 7I; wheat: *P* = 0.050, Figure S6H). No significant differences were evident between aphid strains when considering the total number adults and total number of nymphs, regardless of plant species (*P* > 0.050 in all cases).

### Rickettsiella *impacts alate production and virulence to wheat plants in mesocosms*

As above, virulence to wheat plants in the mesocosm experiment showed patterns consistent with our earlier experiments using individual cages (Figures 4, 5 & S6), where the *Rickettsiella* (+) treatment was shown higher virulence across time compared with the *Rickettsiella* (-) treatment (Figure 8). The most obvious difference between *Rickettsiella* treatments was observed at week 3.5 on the release plants, with fewer leaves (t-test: *P* < 0.001, Figure 8B), a decrease in total leaf area (*P* = 0.050, Figure 8C), a higher aphid infestation score (*P* < 0.001, Figure 8F) and more severe plant damage (*P* = 0.007, Figure 8G) recorded for wheat plants exposed to *Rickettsiella* (+) aphids. Additionally, earlier and greater chlorotic streaking were observed in the *Rickettsiella* (+) infested plants, with a significant difference at week 3 (*P* = 0.046, Figure 8D). For the repeated measurements, wheat plants infested with *Rickettsiella* (+) aphids had fewer leaves (GLM: F_1,38_ = 11.368, *P* = 0.002), a smaller leaf area (F_1,38_ = 14.849, *P* < 0.001), higher chlorotic streaking before damage levels converged (week 2-3.5: F_1,31_ = 7.409, *P* = 0.011) and greater plant damage scores (F_1,38_ = 12.414, *P* = 0.001). When a combination of all plant traits was considered in the MANOVAs, virulence to wheat plants differed significantly between strains at week 2 (*P* = 0.023) and week 3 (*P* = 0.050) but differences were marginally non-significant at week 4 (*P* = 0.070).

**Figure 8.**
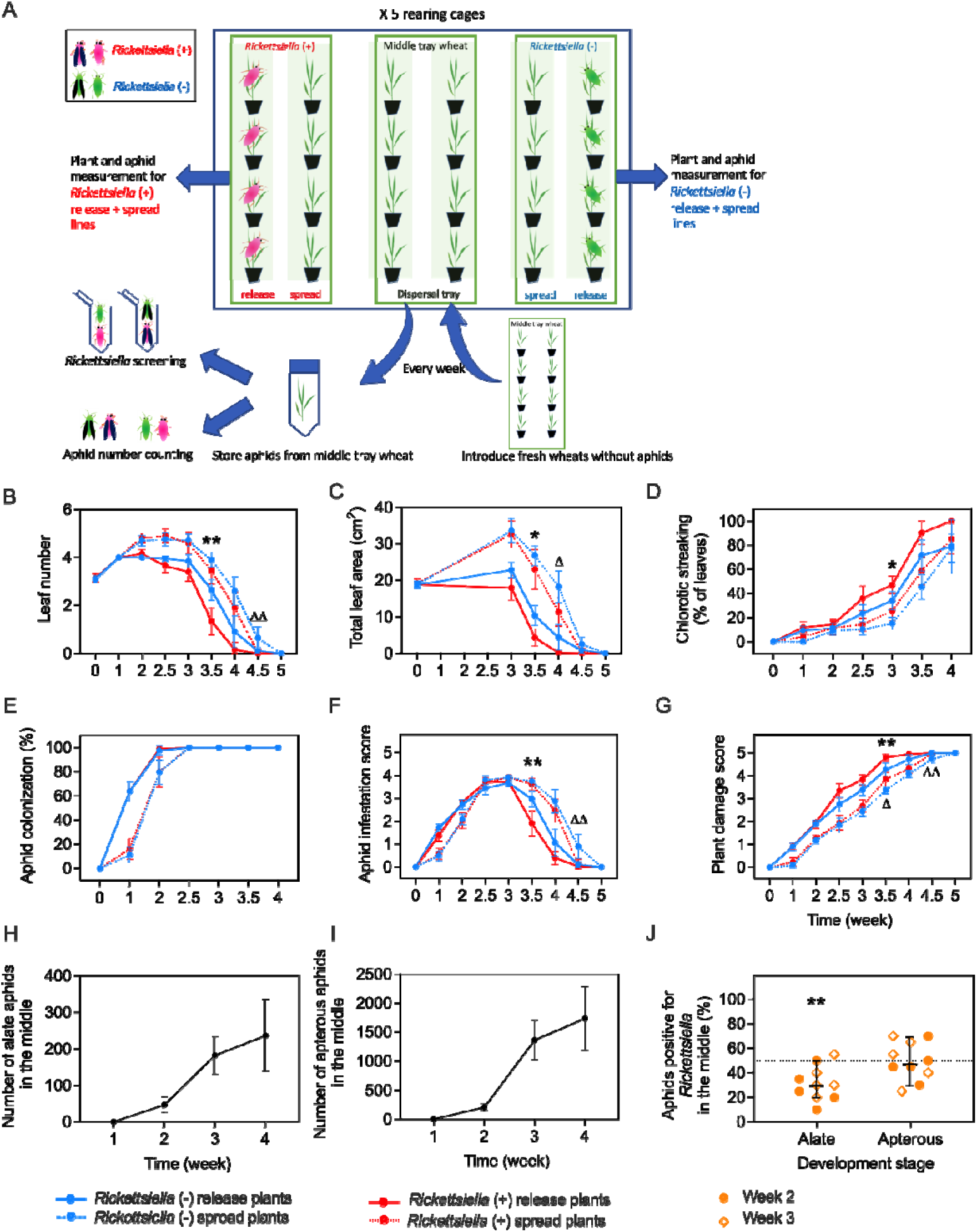
*Rickettsiella* effects on the virulence and dispersal of *D. noxia* on wheat plants in mesocosms. (A) Experimental design. Populations were initiated with 5 *Rickettsiella* (+) or *Rickettsiella* (-) *D. noxia* on wheat plants (“release wheat”) while adjacent plants (“spread wheat”) were monitored. All plants were measured for (B) leaf number, (C) total leaf area, (D) chlorotic streaking, (E) aphid colonization rate, as well as (F) an aphid infestation score and (G) plant damage score. “*” and “**” represent significant differences at *P* < 0.05 and *P* < 0.01, respectively when comparing *Rickettsiella* (+) and *Rickettsiella* (-) on release wheat plants. “Δ” and “ΔΔ” represent significant differences at *P* < 0.05 and *P* < 0.01, respectively when comparing *Rickettsiella* (+) and *Rickettsiella* (-) on spread plants. Aphids collected from the middle tray (“dispersal”) were counted and screened to determine (H) the number of alate adults, (I) the number of apterous adults and (J) the percentage of alate and apterous adults positive for *Rickettsiella*. Symbols in (J) represent the percentage of 20 alate and 20 apterous adults positive for *Rickettsiella* at week 2 and 3. *“**”* represents a significant difference at *P* < 0.01 as determined from a paired t-test.

Increased virulence from *Rickettsiella* (+) aphids was also evident in the spread plants that were placed adjacent to release plants in the mesocosms (Figure 8A). This was particularly evident at week 4.5, with fewer leaves (t-test: *P* = 0.003, Figure 8B), a higher aphid infestation (*P* = 0.001, Figure 8F), and higher plant damage score (*P* = 0.001, Figure 8G) in the *Rickettsiella* (+) treatment compared with the *Rickettsiella* (-) treatment. The total leaf area was also lower in the *Rickettsiella* (+) treatment, with a significant difference at week 4 (*P* = 0.020, Figure 8C). For the repeated measurements taken in the spread plants, once again there was a similar pattern to those in the release plants where plants infested with *Rickettsiella* (+) aphids had a smaller leaf area (F_1,38_ = 4.432, *P* = 0.042) and greater plant damage score (F_1,38_ = 6.009, *P* = 0.019). Despite significant effects on individual plant traits, the damage to wheat plants did not differ significantly among treatments in the spread plants at weeks 2 to 4 (MANOVA: *P* > 0.050 in all cases).

At week 2 and again at week 3, the percentage of alate aphids positive for *Rickettsiella* from the middle tray of plants was significantly below the expected frequency of 50% (t-test: *P* = 0.002, Figure 8J). In contrast, we did not find any significant deviation from 50% for the apterous aphids that were positive for *Rickettsiella* collected from the middle tray (*P* = 0.922). As expected, the number of alate (Figure 8H) and apterous adults (Figure 8I) observed in the middle trays increased with time.

## Discussion

Here we describe the first transfer of an endosymbiont (*Rickettsiella*) into the novel aphid host *D. noxia* and the first case of an endosymbiont directly influencing aphid virulence to host plants. Furthermore, *D. noxia* carrying the *Rickettsiella* showed a decrease in the production of alates, which can impact the short-distance dispersal of these aphids. Our study demonstrates that phenotypes generated by transinfected *Rickettsiella* are species-specific (when compared to *M. persicae*) and highlights novel effects of this endosymbiont on aphid virulence and dispersal.

After its transfer from *A. pisum*, *Rickettsiella* was stable in *D. noxia*, maintaining a frequency of 100% for more than 50 generations in the laboratory. Surprisingly, the presence of *Rickettsiella* had no impact on the primary endosymbiont *Buchnera* and relatively limited influence on aphid fitness. Previous work has shown that *Rickettsiella* modifies body color and can cause large reductions in fecundity in both the native host *A. pisum* (Tsuchida et al., 2010, Tsuchida et al., 2014) and the transinfected host *M. persicae* (Gu et al., 2023). Conversely, we found no effect on body color and identified two traits (development time and heat tolerance measured in two ways) where *Rickettsiella* appears to provide a modest fitness benefits in *D. noxia*. These benefits could help *Rickettsiella* to persist and perhaps spread in wild populations, though we did not identify clear evidence of spread in our laboratory experiments. The contrasting effects of this infection across aphid species suggest substantial host-dependent functional variation of the same infection.

Despite wide-ranging effects of endosymbionts on aphid phenotypes, few studies have considered impacts of endosymbionts on their host plants. In this study, we found the presence of *Rickettsiella* in *D. noxia* increased virulence to wheat at two different plant growth stages (GS13 and GS39) and when using different experimental scales (microcosms with individual wheat plants and mesocosms with multiple plants). Additionally, we found these effects were not limited to wheat, with increased virulence also detected in barley grass and brome grass, two agriculturally important ‘green-bridge’ plants for *D. noxia* (Ward et al., 2020). Effects of *Rickettsiella* were evident for several plant traits across our experiments, though the severity of damage caused by *Rickettsiella* (+) aphids depended on the plant growth stage. While feeding, *D. noxia* injects a toxin into plants causing chlorosis, streaking and rolling leaves in a number of cereal species (Damsteegt et al., 1992). The presence of *Rickettsiella* (+) led to an increase in chlorotic streaking in wheat plants, as well as in barley grass and brome grass - where such symptoms have been rarely observed in the field within Australia (Pirtle et al., 2019).

The mechanism underpinning this increased virulence is unclear but could involve changes in feeding behaviour, metabolism or toxic saliva. Recent work has demonstrated that other endosymbionts can alter probing and feeding behaviour (Leybourne et al., 2020, Li et al., 2023) or the composition of salivary proteins (Loudit et al., 2018) in aphids. Virulent biotypes of *D. noxia* are known to produce more additional energy than less virulent biotypes under stressful conditions based on measures of the NAD+/NADH ratio (De Jager, 2014), and such differences could also be impacted by endosymbionts (Clavé et al., 2022). Luna et al. (2018) found a close interaction between *D. noxia* and bacteria, with aphid saliva containing bacterial proteins that facilitate aphid virulence to wheat. Apart from bacteria, recent studies have also showed non-coding RNA (Chen et al., 2020) and epigenetic effects (du Preez et al., 2020) that might alter virulence to host plants. It would therefore, be interesting to compare salivary toxins between *Rickettsiella* (+) and *Rickettsiella* (-) aphids and relevant epigenetic pathways in a future study. This information could help to elucidate the mechanism(s) used by *D. noxia* to develop new virulent populations despite a lack of genetic diversity (Puterka et al., 2014).

Wing polymorphism is an important adaptative response to aid aphid dispersal, especially under environmental changes (Simon and Peccoud, 2018). Previous studies have shown that the facultative endosymbiont *Serratia* promotes wing development in *A. pisum* to enhance its dispersal (Kang et al., 2022) and suppression of *Buchnera* through antibiotic treatment reduces the formation of winged morphs in *Sitobion avenae* (Zhang et al., 2015). Our results demonstrate potential impacts of *Rickettsiella* on wing development and alate dispersal. In our experiments involving individual plants of wheat, barley grass and brome grass, as well as the mesocosm experiment with trays of wheat plants, *Rickettsiella* (+) aphids had a reduced frequency of alates but not a reduction in the total number of adults or nymphs. This may reflect effects of *Rickettsiella* on the dispersal ability of alates or the frequency at which they are produced, though we did not directly measure the effects of *Rickettsiella* on wing formation as has been done in other studies (e.g. Kang et al., 2022). Our design was limited by our ability to assess aphids present on plants but not in the soil or other areas of the arenas that we used. Given the sensitivity of wing formation to environmental conditions (Müller et al., 2001), additional experiments across multiple generations, and on a larger scale with more capacity for mobility, are needed to establish and the full extent of *Rickettsiella* impacts wing formation and the dispersal of aphid morphs.

Our results highlight the importance of comprehensive assessments of endosymbiont transfers before considering any field application. While *Rickettsiella* may potentially suppress *D. noxia* in the field through a combination of decreased aphid dispersal and increased virulence to ‘green-bridge’ plants that are important in sustaining populations between cropping seasons, further evidence is needed to determine if these phenomena outweigh the fitness advantages and increased virulence to wheat. More genetic clones of *D. noxia* should also be transinfected and tested in the future given endosymbiont effects can vary with host genotype (Lukasik et al., 2013, Niepoth et al., 2018). This work would be challenging to conduct in Australia because only a single genotype has been identified in that country (Yang et al., unpublished data), reflecting the recent incursion of this species from overseas (Yazdani et al., 2018). Additionally, it would also be interesting to test the effects of *Rickettsiella* on more plant hosts of agricultural importance that are attacked by *D. noxia,* such as barley and durum wheat (Hughes, 1990). And finally, further exploration is warranted under greenhouse and/or field conditions, where a host of other factors not testable in laboratory experiments are clearly important.

## Data Availability Statement

All data associated with the article are provided in the manuscript and its supplementary files.

## Acknowledgements

We thank Courtney Brown and Kelly Richardson for technical assistance, the Grains Innovative Park for supplying the *D. noxia* colony and Christopher Preston for providing barley grass and brome grass seeds. This work was undertaken as part of the Australian Grains Pest Innovation Program (AGPIP), supported through funding provided by the Grains Research and Development Corporation (UOM1905-002RTX) and the University of Melbourne.

---

**Figure S1.**
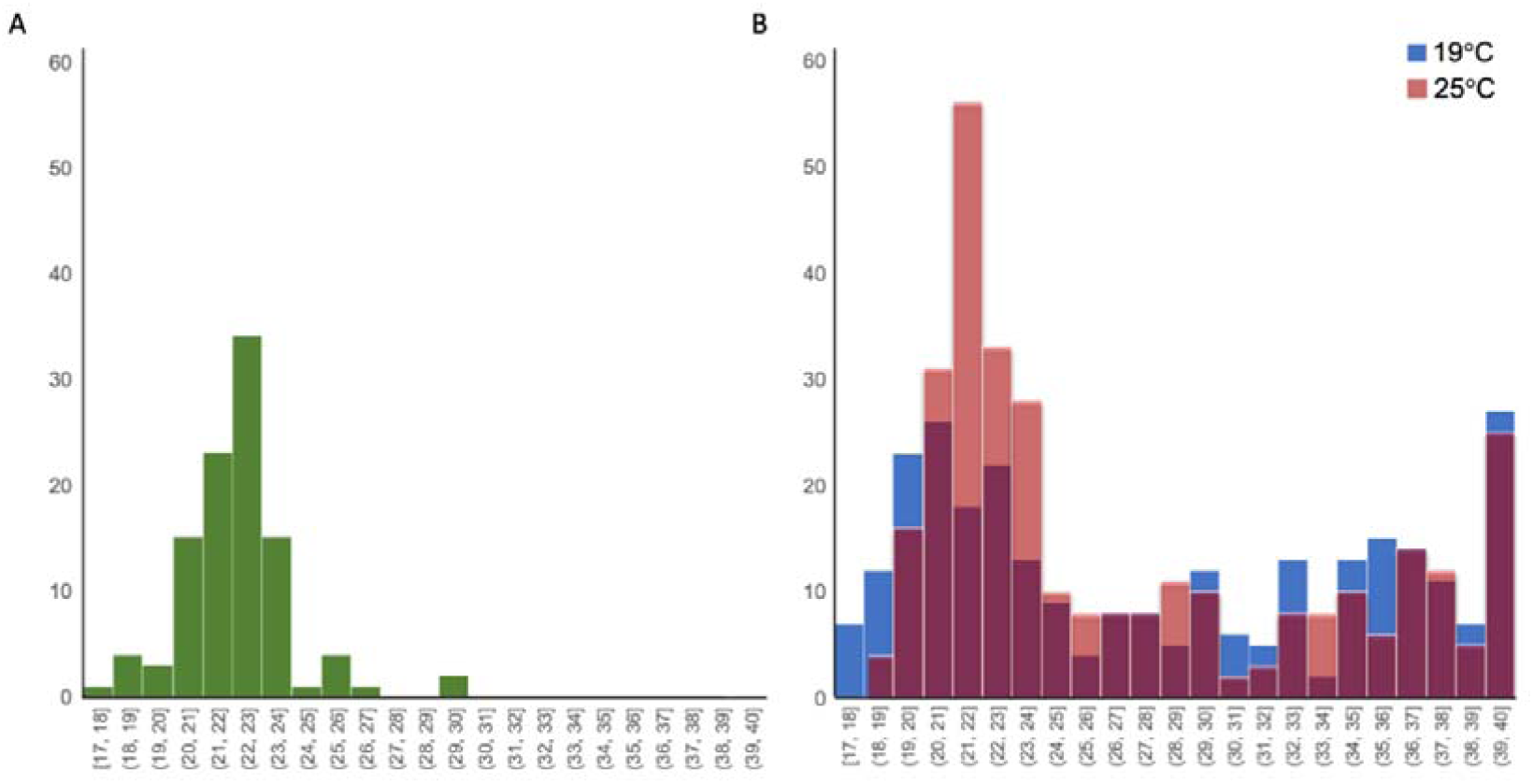
*Rickettsiella* Cp values in (A) laboratory *Rickettsiella* (+) colonies for routine screening and (B) the BugDorm experiment measuring persistence at 19 °C and 25 °C.

**Fig S2.**
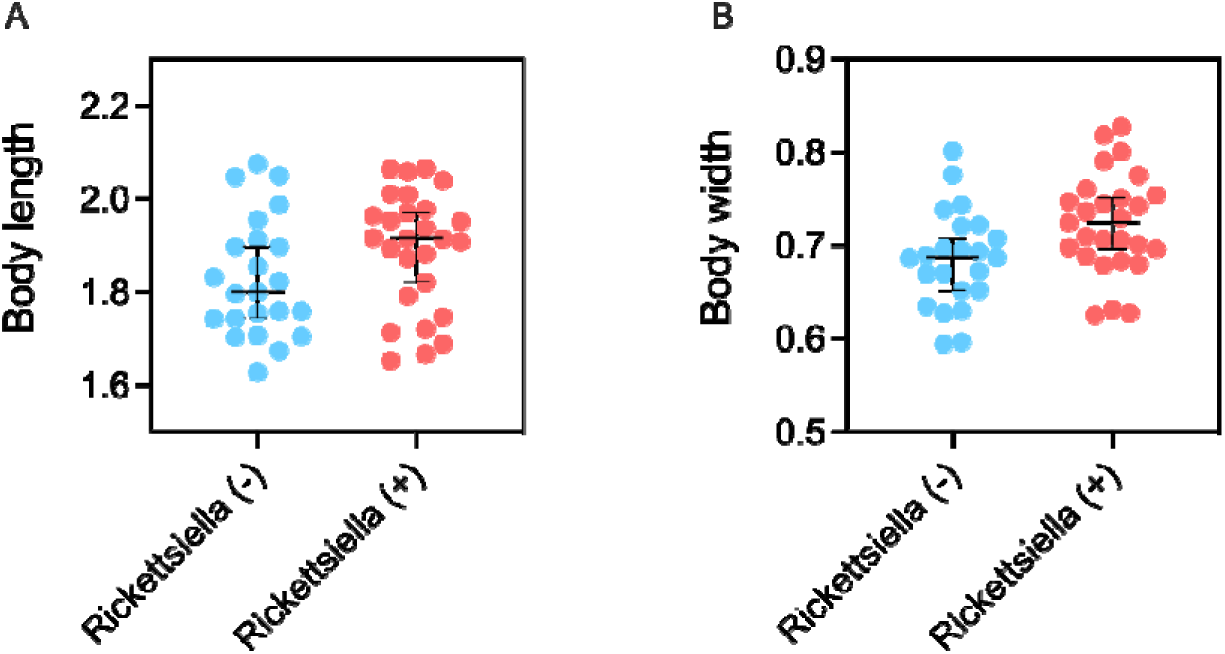
Effect of *Rickettsiella* infection on *D. noxia* (A) body length and (B) body width.

**Figure S3.**
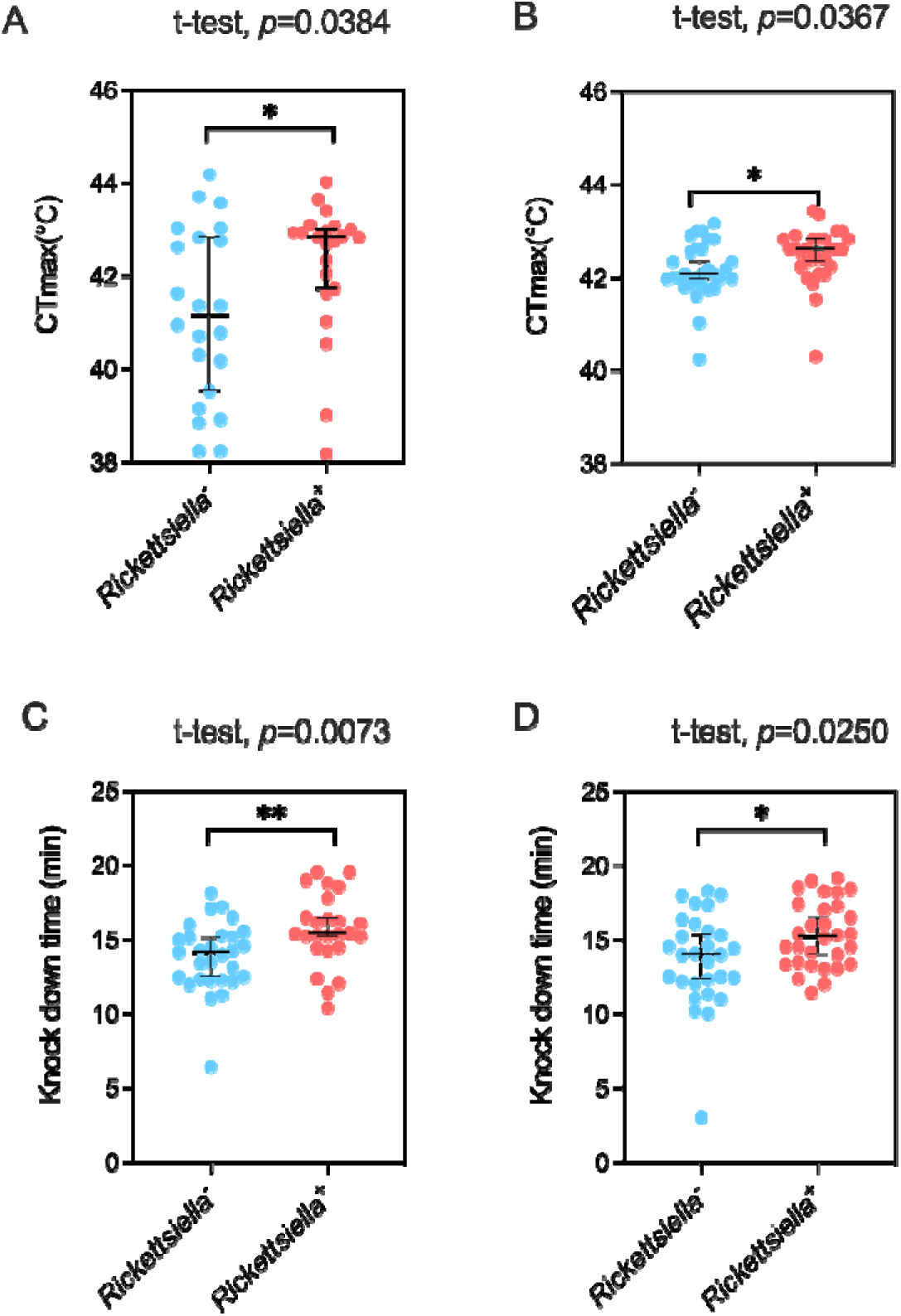
Effect of *Rickettsiella* infection on *D. noxia* heat tolerance. Data is separated by block 1 (A and C) and block 2 (B and D).

**Figure S4.**
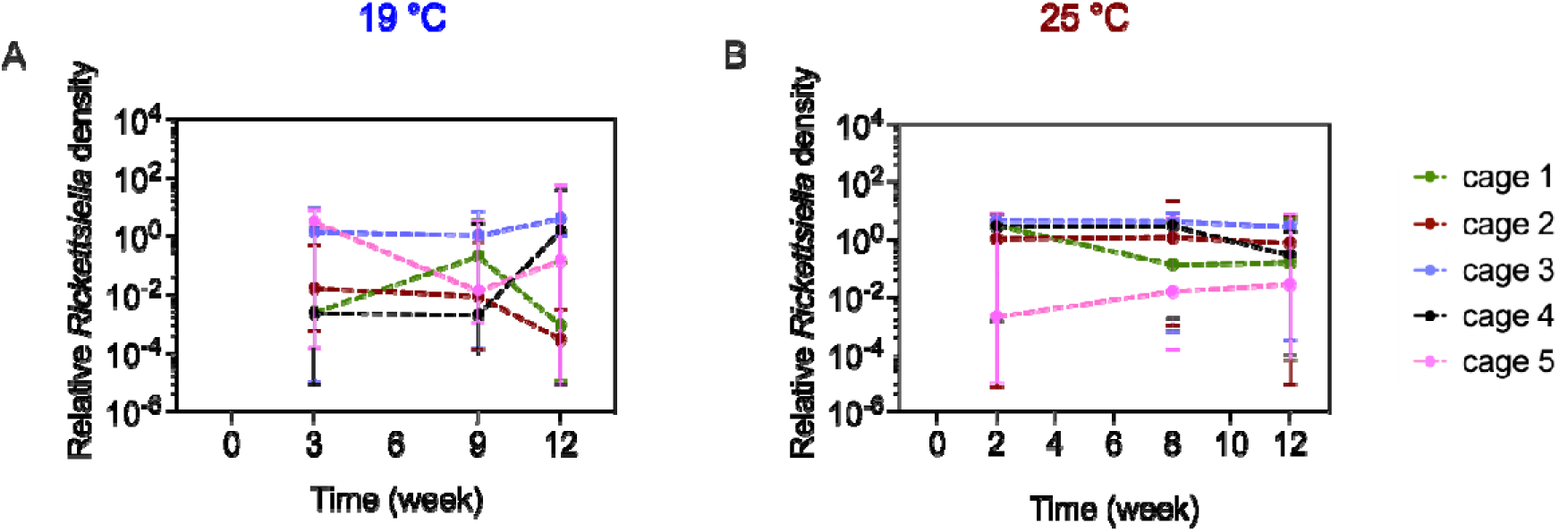
*Rickettsiella* relative density in *D. noxia* mixed population cages at (A) 19 °C and (B) 25 °C.

**Figure S5.**
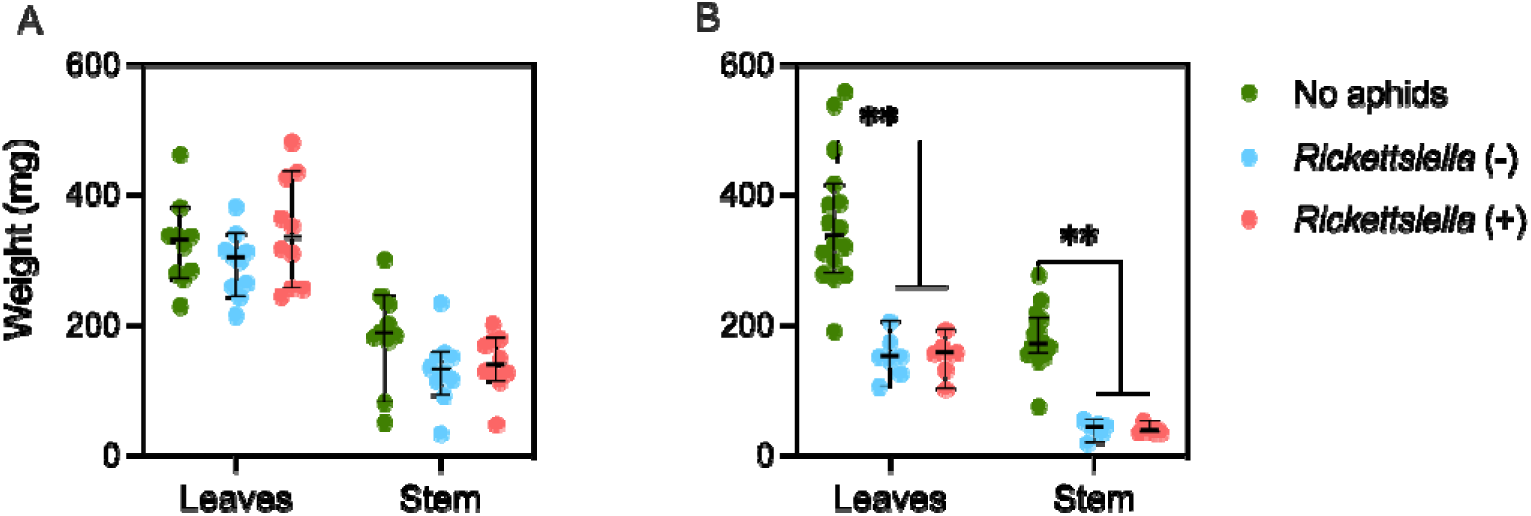
Plant biomass (mg) in (A) GS39 and (B) GS13 wheat plants after the virulence experiments.

**Figure S6.**
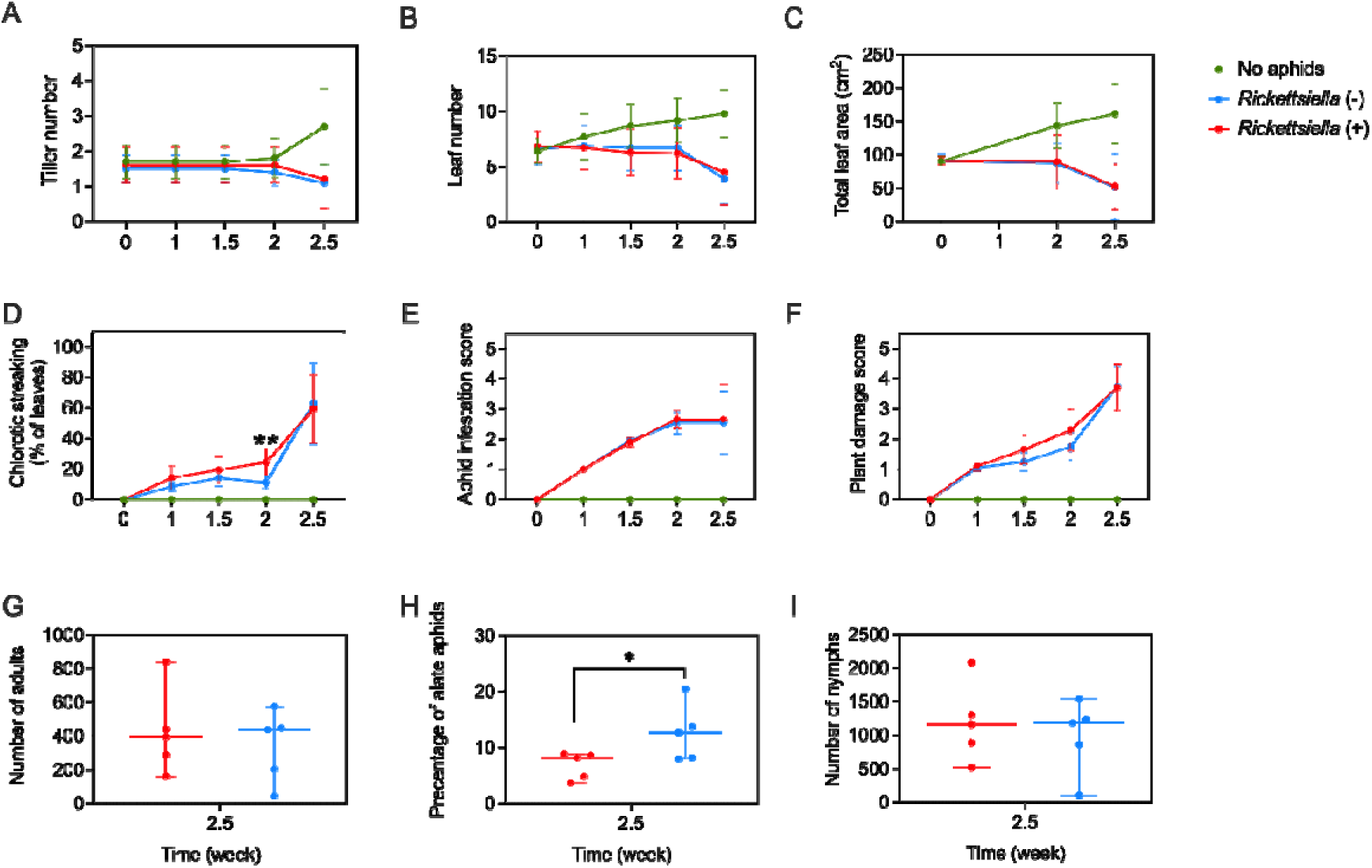
Aphid virulence to wheat plants at GS13 when infested with *Rickettsiella* (+) or *Rickettsiella* (-) *D. noxia* and effects on aphid developmental stage. Three age-matched *Rickettsiella* (+) or *Rickettsiella* (-) *D. noxia* were placed on individual wheat plants. Clean plants without aphids were included for comparison. Each treatment had 5 replicates with 2 plants per replicate and measured traits were (A) tiller number, (B) leaf number, (C) total leaf area, (D) chlorotic, as well as (E) an aphid infestation score and (F) plant damage score. At week 2.5, aphids from the *Rickettsiella* (+) or *Rickettsiella* (-) treatments were removed and the number of alate adults, apterous adults and nymphs recorded. (G) The number of adults (alate plus apterous adults), (H) the percentage of alates and (I) the number of nymphs were used to characterize the developmental stages of aphids. “*” and “**” represent significant differences at *P* < 0.05 and *P* < 0.01, respectively comparing *Rickettsiella* (+) and *Rickettsiella* (-) treatments by independent sample t-tests.

